## Supplementary material for "Spatial summation of pain is associated with pain expectations: Results from a home-based paradigm": Files combined

**Supplementary Material 1. Means and standard deviations for temperatures in pilot study (training)**

| **Time of measurement** | **Segment** | **Ascending (SD)** | **Descending (SD)** |
| --- | --- | --- | --- |
| 10s | 1 | 5.28°C (0.44) | 4.91°C (0.51) |
| 30s | 1 | 5.30°C (0.43) | 4.94°C (0.48) |
| 50s | 1 | 5.33°C (0.40) | 4.91°C (0.51) |
| 10s | 2 | 5.22°C (0.35) | 5.10°C (0.46) |
| 30s | 2 | 5.20°C (0.40) | 5.08°C (0.53) |
| 50s | 2 | 5.24°C (0.41) | 5.14°C (0.43) |
| 10s | 3 | 5.10°C (0.26) | 5.05°C (0.26) |
| 30s | 3 | 5.05°C (0.35) | 5.05°C (0.27) |
| 50s | 3 | 5.04°C (0.32) | 5.08°C (0.25) |
| 10s | 4 | 5.13°C (0.25) | 5.16°C (0.24) |
| 30s | 4 | 5.11°C (0.35) | 5.00°C (0.46) |
| 50s | 4 | 5.01°C (0.36) | 4.94°C (0.50) |
| 10s | 5 | 5.12°C (0.37) | 5.13°C (0.15) |
| 30s | 5 | 4.93°C (0.41) | 5.04°C (0.19) |
| 50s | 5 | 4.89°C (0.43) | 5.03°C (0.27) |

SD - standard deviations

**Supplementary Material 2. Protocol deviations**. The following deviations from the pre-registered protocol must be acknowledged: i) the declared sample of N=90 was not reached due to rigorous inclusion criteria and the peak of incidence rates of COVID-19 during recruitment for this experiment, ii) correlations are reported based on Spearman`s rank correlation coefficient, iii) apart from polynomial contrast results, pairwise comparisons are reported, iv) neither outcomes nor residuals were normally distributed so General Linear Model was replaced by General Estimated Equations, however both analyses provided similar results.

**Supplementary Material 3. Exclusion criteria.** Specific exclusion criteria: Raynaud's phenomenon (cold fading of the fingers, followed by blush numbness and redness), cold allergy (cold urticaria), cold intolerance, cryoglobulinemia, paroxysmal cold hemoglobinuria, rheumatic diseases (e.g. osteoarthritis, rheumatoid arthritis, fibromyalgia, systemic lupus erythematosus, etc.), pheochromocytoma, skin sensitivity disorders, sympathetic neuropathies, cardiovascular diseases (coronary artery disease, chronic heart failure, cardiac insufficiency), neuropathy (e.g. cardiovascular disorders), adrenal pheochromocytoma, sensory skin disorders, sympathetic neuropathies, cardiovascular diseases (coronary artery disease, chronic heart failure class III and IV according to NYHA), hypothyroidism, purulent gangrenous skin lesions, local blood flow disorders.

**Supplementary Material 4. Study procedures. Sequence of conditions was random, “ascending” is presented as an example**

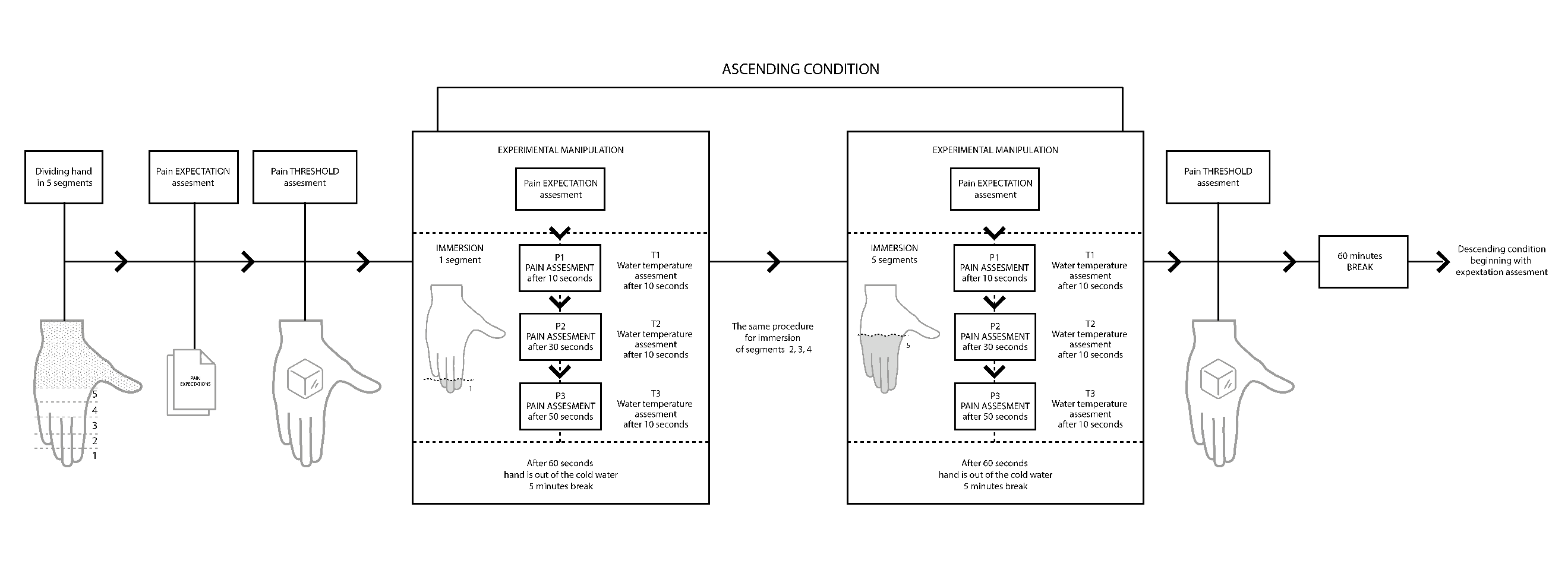

**Study procedures.** In each condition single trial lasted 60 seconds (regardless of the number of segments involved). Each trial started with a question about expected pain intensity. Before and after each experimental condition cold pain thresholds (PT_COLD_) were tested on the examined limb. Participants were instructed to immerse their hand up to the line which separated a given number of segments. Participants were prompted to rate their pain intensity on the VAS scale at the following time points: after 10, 30 and 50s. Inter-trial intervals were set at 5 minutes. The interval between each condition was one hour.

**Supplementary Material 5. Measurement of pain thresholds**

| **Pain threshold (seconds)** | **Mean (SD)** |
| --- | --- |
| Before Ascending | 73.28 (134.88) |
| After Ascending | 76.53 (125.87) |
| Before Descending | 74.10 (139.38) |
| After Descending | 55.73 (85.08) |

**Supplementary Material 6. Means and standard deviations for pain intensity reported during cold water immersions**

| **Condition** | **Time of measurement** | **Segment(s)** | | | | |
| --- | --- | --- | --- | --- | --- | --- |
|  |  | **1/5** | **2/5** | **3/5** | **4/5** | **5/5** |
|  |  | **Mean (SD)** | **Mean (SD)** | **Mean (SD)** | **Mean (SD)** | **Mean (SD)** |
| **Ascending** | **10 s** | 8.54 (11.31) | 10.60 (12.27) | 13.85 (15.18) | 17.44 (17.45) | 22.77 (20.39) |
|  | **30 s** | 13.82 (15.42) | 16.23 (16.39) | 21.13 (20.53) | 27.97 (23.35) | 34.89 (25.43) |
|  | **50 s** | 18.92 (20.09) | 23.07 (20.40) | 30.63 (25.09) | 36.57 (26.53) | 43.19 (29.24) |
| **Descending** | **10 s** | 6.17 (11.49) | 10.64 (14.35) | 14.30 (14.59) | 20.42 (18.63) | 23.48 (21.95) |
|  | **30 s** | 8.50 (13.32) | 13.61 (17.60) | 20.45 (19.84) | 29.73 (24.07) | 36.50 (25.62) |
|  | **50 s** | 10.79 (15.44) | 17.19 (20.62) | 35.63 (23.16) | 37.80 (25.56) | 47.73 (29.10) |

**Supplementary Material 7. Slopes and intercepts for relationships between pain and number of stimulated segments**

|  |  | **Ascending** | | | |  | **Descending** | | | |
| --- | --- | --- | --- | --- | --- | --- | --- | --- | --- | --- |
| **Variables** | **R^2^** | **B** | **SE** | **β** | ***p*** | **R^2^** | **B** | **SE** | **β** | ***p*** |
| 10s | 0.97 | 3.53 | 0.34 | 0.98 | < 0.01 | 0.99 | 4.44 | 0.23 | 0.99 | < 0.001 |
| 30s | 0.96 | 5.38 | 0.55 | 0.98 | < 0.01 | 0.99 | 7.21 | 0.39 | 0.99 | < 0.001 |
| 50s | 0.99 | 6.20 | 0.27 | 0.99 | < 0.001 | 0.98 | 9.45 | 0.60 | 0.99 | < 0.001 |
| Mean | 0.98 | 5.08 | 0.38 | 0.99 | < 0.001 | 0.99 | 7.06 | 0.39 | 0.99 | < 0.001 |

10s - pain intensity measured after 10 seconds, 30s - after 30 seconds, 50s - after 50 seconds of immersion, Mean - mean pain intensity from each immersion, B - unstandardized coefficients, SE - standard error, β - standardized coefficients, *p –* significance value.

**Supplementary Material 8. Pain-related expectations measured prior to cold water immersions**

|  | **Ascending** | | | | **Descending** | | | |
| --- | --- | --- | --- | --- | --- | --- | --- | --- |
| **Segment** | **Mean** | **SD** | **M** | **IQR** | **Mean** | **SD** | **M** | **IQR** |
| 1/5 | 20.77 | 21.91 | 14 | 29 | 22.45 | 21.44 | 16.5 | 29.5 |
| 2/5 | 23.58 | 19.99 | 15 | 29 | 25.70 | 19.38 | 23.5 | 25.5 |
| 3/5 | 25.85 | 19.70 | 21 | 29 | 29.67 | 19.23 | 28 | 28.5 |
| 4/5 | 34.74 | 23.60 | 31 | 37 | 34.27 | 21.09 | 33 | 32 |
| 5/5 | 42.04 | 27.11 | 42 | 51 | 37.05 | 23.58 | 36.5 | 40 |

1/5 – Segment 1, 2/5 – Segments 1 to 2, 3/5- Segments 1 to 3, 4/5 – Segments 1 to 4, 5/5- Segments 1 to 5, SD, standard deviations. M, median. IQR, interquartile range.

**Supplementary Material 9. The number of participants tested by each examiner**

| **Examiner** | **Number of assessed participants** | **Percent** |
| --- | --- | --- |
| E1 | 4 | 5.882 |
| E2 | 4 | 5.882 |
| E3 | 3 | 4.412 |
| E4 | 6 | 8.824 |
| E5 | 7 | 10.294 |
| E6 | 9 | 13.235 |
| E7 | 12 | 17.647 |
| E8 | 5 | 7.353 |
| E9 | 2 | 2.941 |
| E10 | 6 | 8.824 |
| E11 | 2 | 2.941 |
| E12 | 1 | 1.471 |
| E13 | 7 | 10.294 |
